# Translational lipidomics reveals BMP and its precursor LPG as biomarkers for CLN5 Batten disease

**DOI:** 10.64898/2026.03.19.712969

**Authors:** Eshaan S. Rawat, Nick Manfred, Hisham N. Alsohybe, Wentao Dong, Isabelle V. Peña, Maryam Idris, Christian Posern, Lena M. Westermann, Samantha J. Murray, Daan M. Panneman, Stefania Della Vecchia, Stefano Doccini, Maria Marchese, Filippo M. Santorelli, Peter M. van Hasselt, Nadia L. Mitchell, Angela Schulz, Monther Abu-Remaileh

## Abstract

CLN5 Batten disease, caused by biallelic mutations in *CLN5*, is a rare, early-onset neurodegenerative lysosomal storage disorder that has no cure and lacks validated biomarkers, hindering accurate diagnosis and assessment of therapeutic response. We recently identified CLN5 as the synthase of bis(monoacylglycero)phosphate (BMP), an endolysosomal phospholipid crucial for lysosome function and lipid catabolism. This suggested BMP and its precursor lysophosphatidylglycerol (LPG) as clinically relevant biomarkers. It also prompted *in vivo* confirmation of CLN5 as the biologically relevant lysosomal BMP synthase. Here we show that murine and ovine disease models lacking CLN5 show significant and universal depletion of BMP and elevation of LPG across tissues and brain regions, consistent with the biochemical function of CLN5. Additionally, lysosomal lysates from murine models of CLN5 Batten disease lack the ability to synthesize BMP from its precursor LPG, establishing CLN5 as the main BMP synthase *in vivo*. Of importance, CLN5 patient-derived fibroblasts show BMP depletion and LPG elevation. Translating these results towards clinical utility, we demonstrate BMP and LPG to be accessible biomarkers for CLN5 Batten disease in both plasma and dried blood spots, enabling early diagnosis and patient screening.

## Introduction

Lysosomes degrade macromolecules and faulty organelles to recycle their content and engage in signaling and metabolism to maintain cellular homeostasis^1–3^. Underscoring their essential roles, monoallelic mutations in lysosomal genes are common risk factors for late-onset neurodegenerative diseases, such as Alzheimer’s disease, Parkinson’s disease, amyotrophic lateral sclerosis, and frontotemporal dementia^4,5^. Biallelic mutations in these genes most often exacerbate dysfunction, causing lysosomal storage disorders (LSDs). One class of LSDs is neuronal ceroid lipofuscinosis (NCL), commonly referred to as Batten disease. NCLs are caused by mutations in one of the *Ceroid Lipofuscinosis Neuronal* (*CLN*) genes, and were initially described based on the pathologic accumulation of the aging-related autofluorescent storage material lipofuscin^6,7^. NCLs are characterized by a heterogeneous combination of childhood dementia, epilepsy, motor decline, seizures, visual failure, progressive cognitive deterioration, and limited life expectancy^8,9^. Several CLN proteins still have unknown or incompletely characterized functions, thus challenging the development of therapeutic interventions. One such functionally elusive protein, until recently, was CLN5, the dysfunction of which is the cause of CLN5 Batten disease, also referred to as variant late infantile NCL (vLINCL) or Finnish variant late infantile neuronal ceroid lipofuscinosis^10,11^. CLN5 Batten disease pathology in the brain includes substantial atrophy of the cerebrum and cerebellum, thinner cerebral cortex, enlarged ventricles, and reduction of white matter, as well as neurodegeneration in the thalamus and striatum^10,12^. Granule cells and Purkinje cells of the cerebellum are particularly susceptible to neuronal atrophy in advanced stages of disease progression, as are layer III and V neurons of the cerebral cortex and pyramidal cells and interneurons in the hippocampus, while retinal dystrophy and degeneration occur at earlier stages^10^.

Work with *Cln5* deficient (*Cln5*^−/−^) mice has correlated phenotypic observations in the murine model with clinical observations of CLN5 Batten disease patients, including accumulation of autofluorescent storage material, regional atrophy, vision loss and retinal degeneration, and defective myelination^13–18^. Similarly, a recently developed zebrafish *cln5*^−/−^ model has replicated key features of the human disease, including locomotor impairment, brain cell apoptosis, lipid accumulation, vision loss, and disrupted calcium homeostasis^19,20^. The most clinically relevant work has been with ovine models, given the many advantages of sheep as a model organism for the human brain and human neurological disorders^21,22^. A naturally occurring ovine model of CLN5 Batten disease has been found to replicate the key features of the human disease^23–27^. In hopes of establishing a novel genetic therapy, adeno-associated virus serotype 9 (AAV9) vectors that encode *CLN5* have been delivered intracerebroventricularly and intravitreally, at variable doses and disease stages^28–30^. While intracerebroventricularly delivered therapy was originally insufficient to correct vision loss, it successfully increased lifespan, delayed the progression of disease, and slowed brain atrophy in the majority of treated sheep, notably tripling the expected lifespan to 60 months in one case administered a high AAV9 dose^30^. In a subsequent iteration of treatment, dual intracerebroventricular and intravitreal *CLN5* gene therapy administration proved to have greater efficacy, successfully attenuating intracranial volume loss and sustaining visual function in both early and advanced stages of disease progression^31^. These promising results have subsequently led to the first in-human Phase I/II clinical trial for CLN5 Batten disease, testing dual intracerebroventricular and intravitreal administration of self-complementary AAV9 (scAAV9) vectors that encode human *CLN5*^31^. These developments prompt further characterization of CLN5 functional roles in innate biological contexts to establish reliable biomarkers that help clinicians diagnose, monitor, and ultimately treat CLN5 Batten disease as treatment options emerge.

Using *in cellulo* models of CLN5 Batten disease and enzymatic activity assays, our lab recently identified CLN5 as the synthase of bis(monoacylglycero)phosphate (BMP), an endolysosomal phospholipid crucial for lysosome structure and its lipid catabolism function^32–34^. Loss of functional CLN5 in HEK293T cells, induced pluripotent stem cells (iPSCs), and iNeurons not only decreased BMP species globally, but also increased the abundance of its direct precursor lysophosphatidylglycerol (LPG)^32,33^. As the primary component of intralysosomal vesicles, BMP is believed to imbue these vesicles with a negatively charged surface that electrostatically recruits lipid hydrolases and potentially activates them in the acidic environment of the lysosome, thereby enabling lipid catabolism^33–38^. BMP dysregulation has been implicated in a number of diseases, including CLN3 Batten disease^39–41^, Gaucher’s disease^42,43^, mixed dementia^44^, and Alzheimer’s disease^45^, as well as with Alzheimer’s disease-associated gene CTSD^46^ and frontotemporal dementia-associated gene GRN^47,48^. Further, BMP has been developed as a known clinical biomarker for drug-induced phospholipidosis, highlighting its clinical relevance and utility^49–51^. BMP has also been shown to be dysregulated in the urine of patients with Parkinson’s disease^52–54^, and has thus been used as a biomarker in LRRK2 inhibitor clinical trials for Parkinson’s disease^55,56^. Beyond BMP, lipids have extensively been studied as biomarkers for a large number of diseases, and have been measured using matrices such as plasma^57–60^ and dried blood spots^61–64^. Specifically, plasma and lipid biomarkers have been studied as biomarkers for diseases including Alzheimer’s disease^65–67^, Parkinson’s disease^68–71^, various cancers^72–78^, and multiple LSDs^79–85^. The proven utility of BMP and other lipids as disease biomarkers suggests BMP and LPG may serve as relevant clinical biomarkers for diagnosing and monitoring CLN5 Batten disease, an incurable disease currently lacking validated biomarkers.

To establish the necessity of CLN5 for BMP synthesis *in vivo* and to address the need for functional biomarker validation, we performed lipidomics across mammalian models of CLN5 Batten disease as well as in samples from CLN5 Batten disease patients, leveraging robust untargeted and targeted LC/MS methods for the detection, annotation, and quantitation of BMP^86^. Murine models of CLN5 Batten disease showed significant depletion of BMP and elevation of LPG at both the whole-tissue and lysosome level across organs. Enzyme assays with lysosomal lysates isolated from these murine models notably required CLN5 to synthesize BMP from LPG in a pH-dependent manner, and were rescued by addition of recombinant CLN5 protein. Ovine models of CLN5 Batten disease similarly showed significant depletion of BMP and elevation of LPG in several brain regions and in the liver. In CLN5 Batten disease children, skin cell-derived fibroblasts, plasma, and dried blood spots showed significantly decreased BMP and increased LPG, suggesting clinical utility of these lipids as biomarkers. Altogether, these results establish CLN5 as the physiological lysosomal BMP synthase by confirming this function in an *in vivo* context, and establish depleted BMP and elevated LPG as accessible biomarkers for CLN5 Batten disease.

## Results

### Cln5^−/−^ mice have significantly depleted BMPs and elevated LPGs across tissues

To test if CLN5 is required for BMP synthesis *in vivo* and assess its impact on lipid metabolism, we utilized a previously developed *Cln5* knockout (*Cln5*^−/−^) mouse line^13^. Whole-tissue untargeted lipidomics on five mouse organs (brain, heart, kidney, liver, and lung) from *Cln5*^−/−^ mice and their *Cln5*^+/−^ control littermates revealed significant depletion of BMP and accumulation of LPG across tissues in *Cln5*^−/−^ mice (Fig. 1A-E and Table S1). Targeted quantitation of BMP and LPG species confirmed these results (Fig. 1F-J), and showed minimal to no change in phosphatidylglycerol (PG), an isomer of BMP (Fig. S1A-E). Of note, for accurate and confident annotation of BMP species, given the challenge of annotating BMP against its isomeric lipid class PG^86^, we established a standardized systematic nomenclature for untargeted Orbitrap LC/MS annotation of BMPs and PGs with three confidence levels (cl1, 2, and 3, with cl1 as the highest, see methods for details). Across all tested tissues, BMP depletion fold changes ranged from one to five orders of magnitude, with a large majority of BMPs having fold changes lower than 1×10^-2^-fold, or even completely undetected in *Cln5*^−/−^ mice (Fig. 1F-J). *Cln5*^+/−^ mice were used as control littermates following initial testing that revealed no significant difference in the levels of BMP, LPG, and PG between *Cln5*^+/+^ mice and *Cln5*^+/−^ mice across tissues (Fig. S2A-O), consistent with Batten disease as an autosomal recessive disorder. Since BMP is a key endolysosomal lipid^32–34^, we further validated our results by profiling lysosome-specific lipidomic changes. Accordingly, we bred *TMEM192-3xHA* (LysoTag) mice^39^ with *Cln5*^−/−^ mice to produce LysoTag-harboring *Cln5*^−/−^ and *Cln5*^+/−^ mice. Lysosome immunoprecipitation (LysoIP) coupled with untargeted lipidomics from brain and liver showed a clear BMP phenotype (Fig. 2A-B and Table S1). Targeted analysis showed that the significant depletion of BMP observed in the lysosome is comparable in fold change to that in bulk tissue lipidomics; fold changes ranged from 3×10^-3^ to 2×10^-4^-fold, with the majority of BMP species completely undetectable in *Cln5*^−/−^ lysosomes (Fig. 2C-D). Notably, LPG accumulation in lysosomes was markedly greater than seen in the whole-tissue. While the maximum LPG fold change in whole-tissues was 26-fold, a large majority of LPG fold changes in lysosomes were over 26-fold, ranging from 14 to 270-fold (Fig. 2C-D). Higher lysosomal LPG accumulation in lysosomal fractions likely reflects direct substrate accumulation from *Cln5* knockout, while slightly lower whole-tissue LPG accumulation may be due to masking of the lysosome-level effect by other pools of extra-lysosomal LPG. As in whole-tissue, lysosomal PG levels exhibited no to minimal changes across tissues (Fig. S1F-G). These findings suggest that CLN5 is necessary for BMP synthesis *in vivo*.

**Figure 1.**
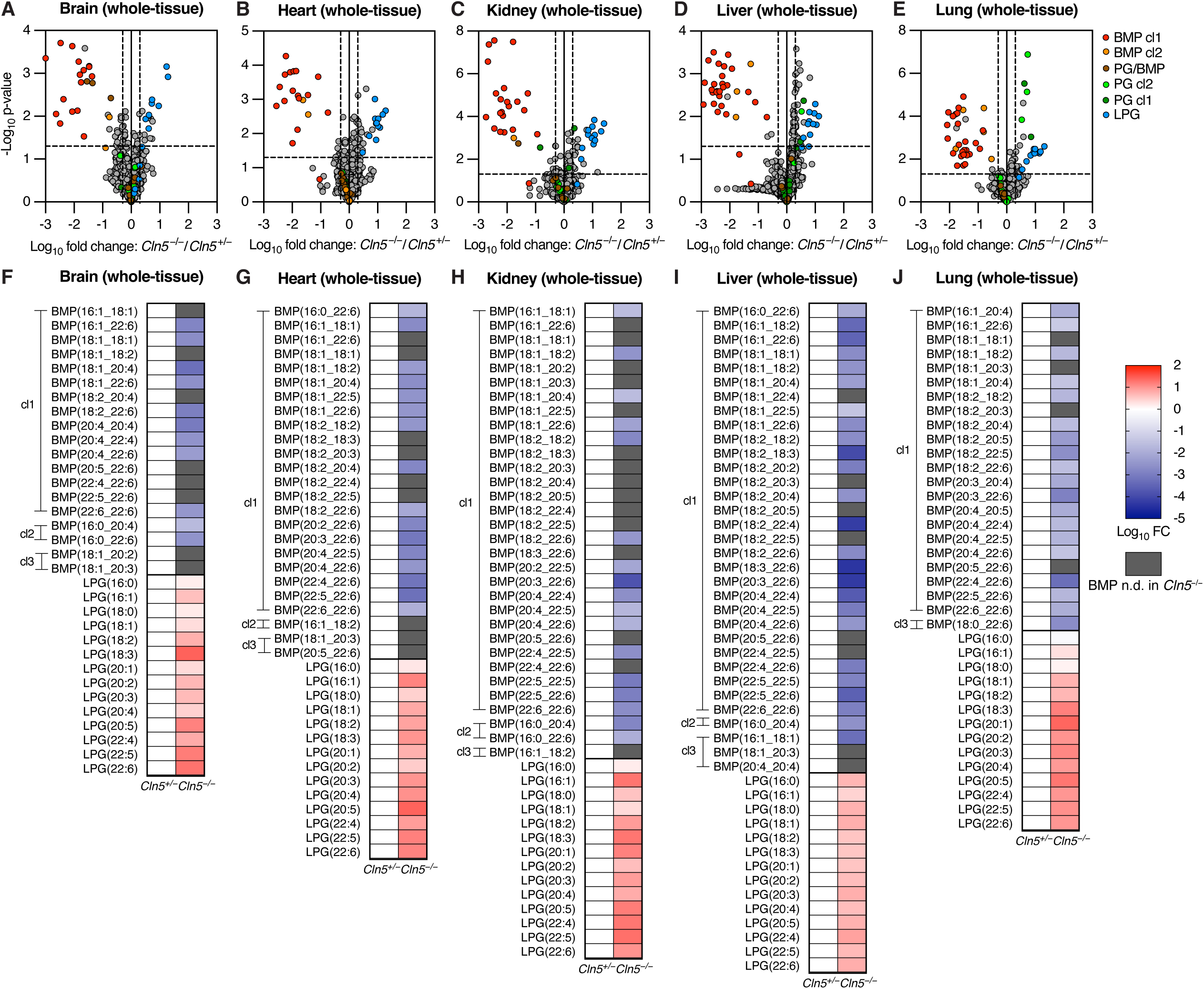
*Cln5*^−/−^ mice exhibit significant depletion of BMPs and accumulation of LPGs across tissues. **A-E**. Volcano plots from untargeted whole-tissue lipidomics of *Cln5*^+/−^ and *Cln5*^−/−^ mice: brain (A), heart (B), kidney (C), liver (D), and lung (E). *Cln5*^−/−^ mice n = 5, *Cln5*^+/−^ mice n = 10 for A-D, *Cln5*^+/−^ mice n = 9 for E. Data from untargeted Orbitrap LC/MS. Fold change (FC) by ratio of means. p-values by Student’s t-tests. Untargeted quantitation volcano plots normalized by constant median normalization. Volcano plot vertical dotted lines presented at FC = 0.5x and 2x (log10 FC = −0.301 and 0.301). Volcano plot horizontal dotted line presented at p-value = 0.05 (−log_10_ p-value = 1.301). **F-J.** Targeted quantitation heatmaps of BMPs and LPGs from untargeted whole-tissue lipidomics, corresponding to above volcano plots: brain (F), heart (G), kidney (H), liver (I), and lung (J). Data from untargeted Orbitrap LC/MS. Fold change (FC) by ratio of means.Targeted quantitation heatmaps normalized by PE(18:1_18:1). Heatmap gray coloring indicates BMP not detected (n.d.) in *Cln5*^−/−^. All highlighted species in volcano plots and quantified species in heatmaps were manually and rigorously validated for proper annotation and integration. BMPs and PGs with confidence level 1 (cl1) label are annotated by definitive +NH_4_ MS2 and −H MS2. BMPs and PGs with cl2 label are annotated by −H MS2 and relative retention time (RT) to an isomeric cl1 PG or BMP in the same tissue. PG/BMP is labeled for species lacking cl1 or cl2, in the volcano plots only. BMPs with cl3 label are annotated by −H MS2 and relative RT to a corresponding cl1 BMP or isomeric PG in a different tissue, in the heatmaps only.

**Figure 2.**
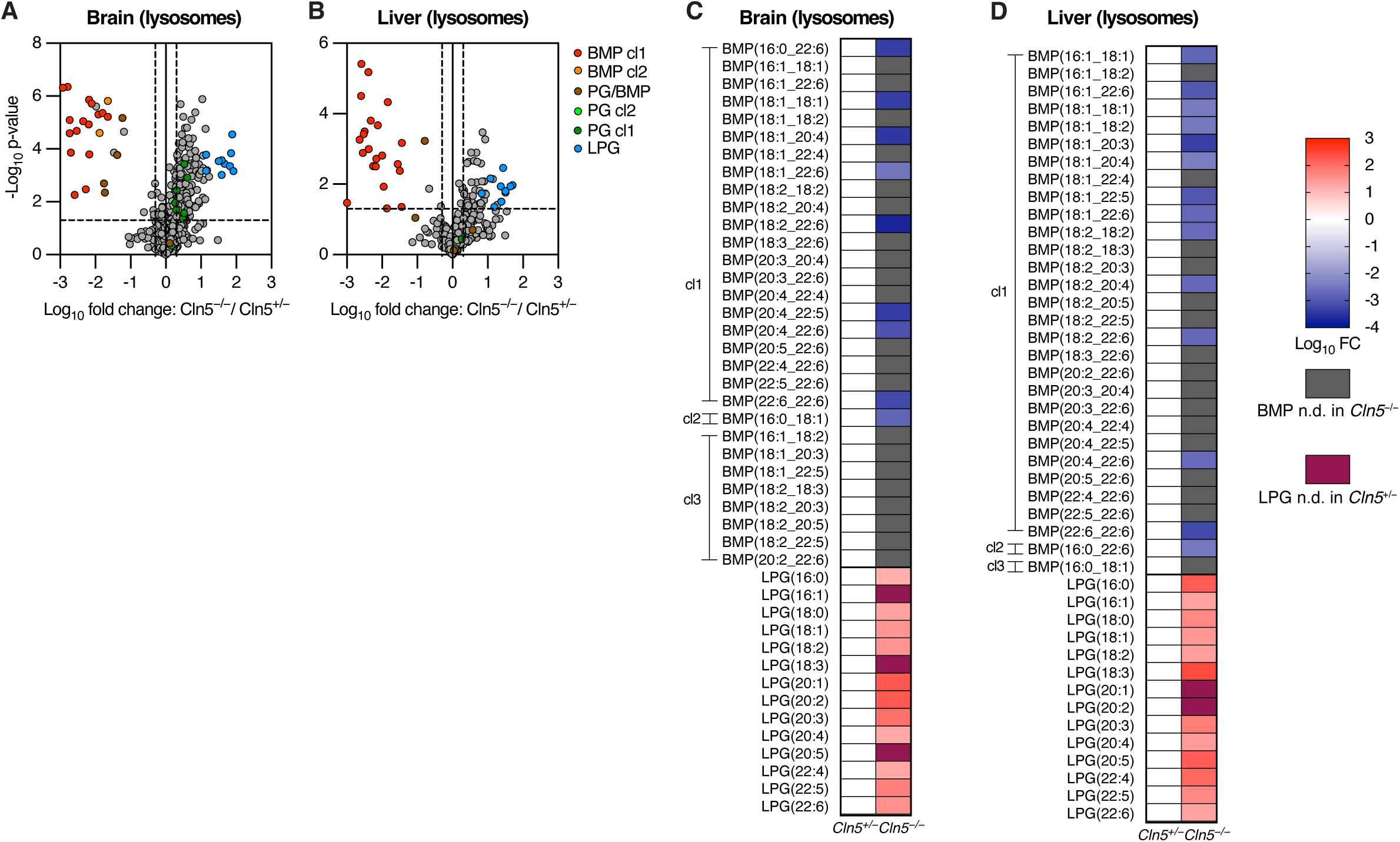
*Cln5*^−/−^ mice lysosomes exhibit significant depletion of BMPs and accumulation of LPGs. **A-B**. Volcano plots from untargeted LysoIP lipidomics of *Cln5*^+/−^ and *Cln5*^−/−^ mice: brain (A) and liver (B). *Cln5*^+/−^ mice n = 4, *Cln5*^−/−^ mice n = 3. Data from untargeted Orbitrap LC/MS. Fold change (FC) by ratio of means. p-values by Student’s t-tests. Untargeted quantitation volcano plots normalized by constant median normalization. Volcano plot vertical dotted lines presented at FC = 0.5x and 2x (log10 FC = −0.301 and 0.301). Volcano plot horizontal dotted line presented at p-value = 0.05 (−log_10_ p-value = 1.301). **C-D.** Targeted quantitation heatmaps of BMPs and LPGs from untargeted LysoIP lipidomics, corresponding to volcano plots: brain (C) and liver (D). Data from untargeted Orbitrap LC/MS. Fold change (FC) by ratio of means. Targeted quantitation heatmaps normalized by PE(18:1_18:1). Heatmap gray coloring indicates BMP not detected (n.d.) in *Cln5*^−/−^. Heatmap purple coloring indicates LPG not detected in *Cln5*^+/−^. All highlighted species in volcano plots and quantified species in heatmaps were manually and rigorously validated for proper annotation and integration. BMPs and PGs with confidence level 1 (cl1) label are annotated by definitive +NH_4_ MS2 and −H MS2. BMPs and PGs with cl2 label are annotated by −H MS2 and relative retention time (RT) to an isomeric cl1 PG or BMP in the same tissue. PG/BMP is labeled for species lacking cl1 or cl2, in the volcano plots only. BMPs with cl3 label are annotated by −H MS2 and relative RT to a corresponding cl1 BMP or isomeric PG in a different tissue, in the heatmaps only.

### CLN5 is required for brain and liver lysosomes to synthesize BMP from LPG

We have previously shown that CLN5 is required for lysosomes derived from HEK293T cells to synthesize BMP from LPG^32^. These observations in isolation do not preclude the existence of potential alternative lysosomal synthases of BMP *in vivo*, prompting us to directly test whether CLN5 is necessary and sufficient for BMP synthesis *in vivo*. To this end, we conducted LysoIP on LysoTag-harboring *Cln5*^+/−^ and *Cln5*^−/−^ mouse brain and liver tissues to isolate lysosomal lysates. We then incubated the lysosomal lysates with deuterated (d5)-LPG(17:0), deuterated on the glycerol adjacent to the 17:0 acyl chain, at a pH of either 5.5 (lysosomal milieu) or 7.4 (cytosolic milieu) (Fig. 3A). In both brain– and liver-derived lysosomal lysates, only *Cln5*^+/−^-derived lysosomal lysates incubated at a pH of 5.5 showed significant synthesis of d5-BMP (Fig. 3B-C). In all other conditions, we were unable to detect d5-BMP significantly above noise levels (Fig. 3B-C). This suggests that BMP synthesis is contingent on a sufficiently acidic environment and that CLN5 is necessary for BMP synthesis. To determine whether CLN5 is sufficient for BMP synthesis, we incubated liver-derived lysosomal lysates from *Cln5*^+/−^ and *Cln5*^−/−^ mice at a pH of 5.5 with d5-LPG and supplemented recombinant CLN5 (rCLN5) protein to *Cln5*^−/−^-derived lysosomes. The rCLN5 addition significantly increased d5-BMP synthesis in a dose-dependent manner relative to *Cln5*^−/−^-derived lysosomes with no rCLN5 added (Fig. 3D). We additionally analyzed d5-BMP production from whole-tissue liver homogenates of *Cln5*^+/−^ and *Cln5*^−/−^ mice incubated with d5-LPG, under pH = 5.5 and pH = 7.4 conditions. Of these four conditions, only *Cln5*^+/−^-derived whole-tissue in a pH of 5.5 resulted in d5-BMP synthesis (Fig. 3E). Thus, CLN5 is necessary and sufficient to synthesize BMP from LPG under physiological conditions and requires a sufficiently acidic milieu.

**Figure 3.**
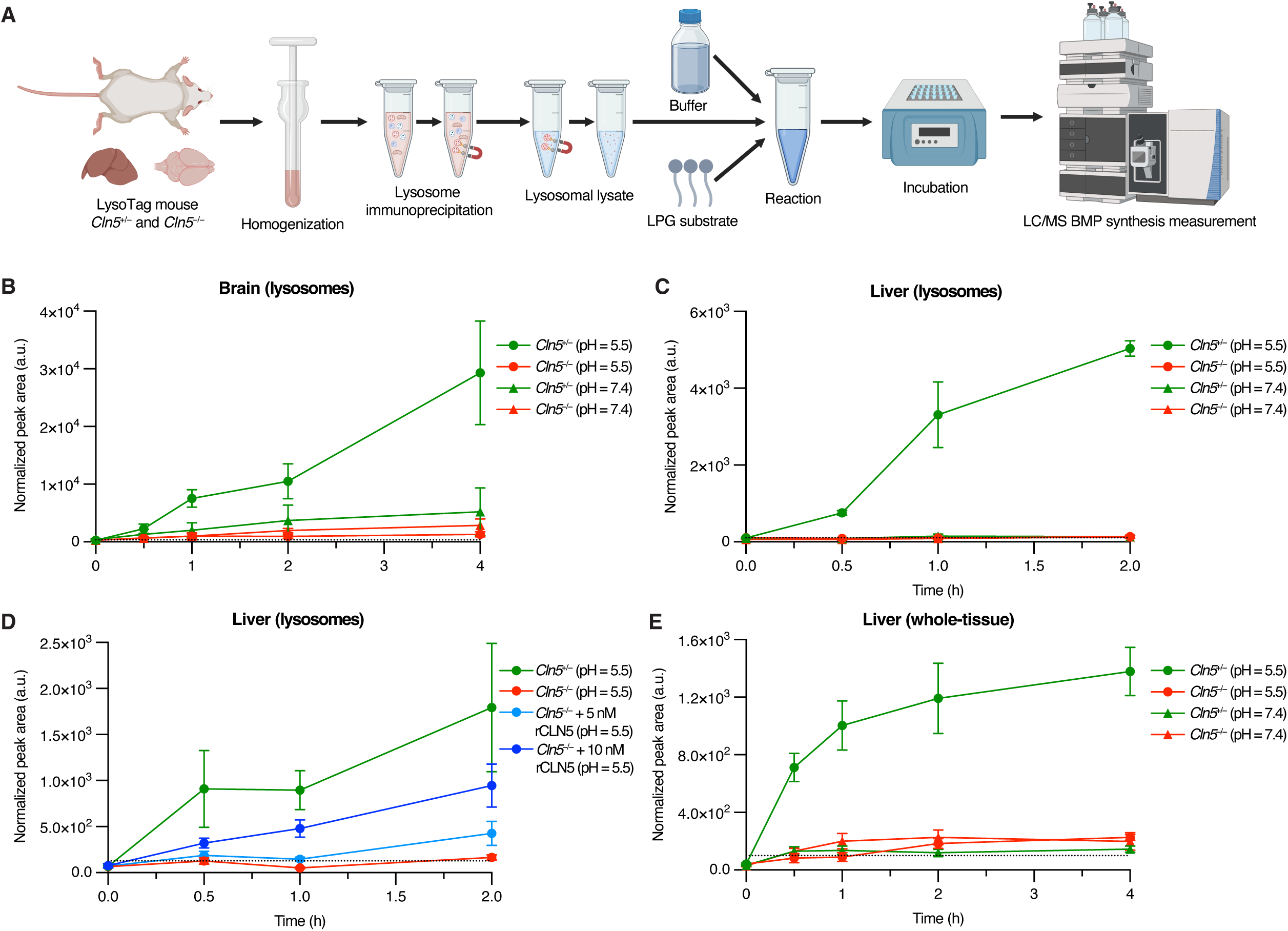
CLN5 is required *in vivo* to synthesize BMP from its precursor LPG. **A.** Depiction of experimental design. **B.** Enzymatic synthesis of d5-BMP from d5-LPG with lysosomal lysates from *Cln5*^+/−^ and *Cln5*^−/−^ mouse brains in acidic and neutral pH. *Cln5*^+/−^ n = 4, *Cln5*^−/−^ n = 4. **C.** Enzymatic synthesis of d5-BMP from d5-LPG with lysosomal lysates from *Cln5*^+/−^ and Cln5^−/−^ mouse livers in acidic and neutral pH. *Cln5*^+/−^ n = 4, *Cln5*^−/−^ n = 4. **D.** Enzymatic synthesis of d5-BMP from d5-LPG with lysosomal lysates from *Cln5*^+/−^ and *Cln5*^−/−^ mouse livers with addition of recombinant CLN5 (rCLN5) in acidic pH as a positive control for the reaction. *Cln5*^+/−^ n = 4, *Cln5*^−/−^ n = 4, *Cln5*^−/−^ + 5 nM rCLN5 n = 4, *Cln5*^−/−^ + 10 nM rCLN5 n = 4. **E.** Enzymatic synthesis of d5-BMP from d5-LPG with whole-tissue lysates from *Cln5*^+/−^ and *Cln5*^−/−^ mouse livers in acidic and neutral pH. *Cln5*^+/−^ n = 5, *Cln5*^−/−^ n = 5. Data from targeted QQQ LC/MS. Data plotted as mean ± SEM. Peak areas normalized by PC(18:1_18:1). Horizontal dotted line in each plot represents the maximum background peak area from all negative control and blank samples.

### CLN5^−/−^ sheep have significantly depleted BMPs and elevated LPGs across brain regions and in the liver

Ovine models of CLN5 Batten disease have been extensively characterized and used as highly relevant models to study disease pathology and preclinically assess CLN5 gene therapy^23–31^. Given the necessity of CLN5 in synthesizing BMP from LPG, and the emerging pattern of BMP depletion and LPG accumulation in cells and mice, we sought to validate this pattern in sheep^21,22^. Using targeted lipidomics, we measured BMP and LPG levels in five distinct brain regions (primary motor cortex, primary visual cortex, thalamus, cerebellum, and parieto-occipital cortex) and liver. Consistent with our previous results, we found that for every examined tissue, *CLN5*^−/−^ sheep have significantly elevated LPG and depleted BMP levels compared to the corresponding region from *CLN5*^+/−^ and *CLN5*^+/+^ sheep (Fig. 4A-F). Analysis of PG levels in these same brain regions and liver again revealed minimal differences in abundance in each region, suggesting moderate relevance, if any, of PG levels to CLN5 Batten disease pathology (Fig. S3A-F). These results indicate that CLN5 is the major BMP synthase across species and establish BMP and LPG as potential biomarkers in relevant preclinical models.

**Figure 4.**
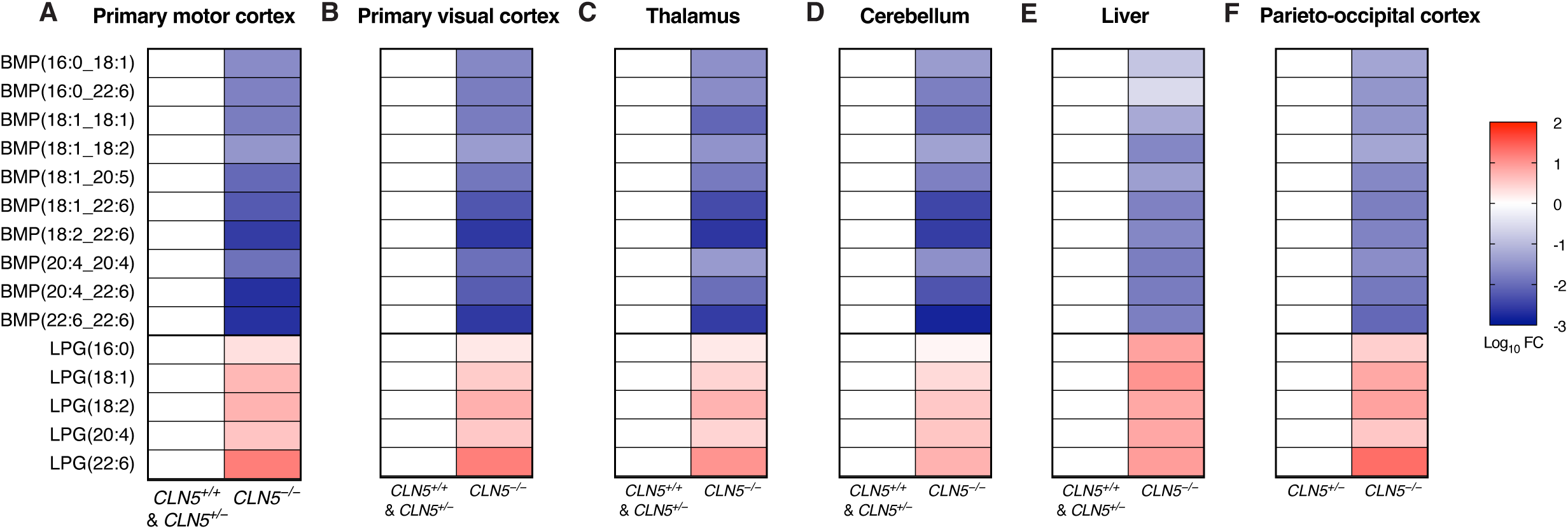
*CLN5*^−/−^ sheep exhibit significant depletion of BMPs and accumulation of LPGs across brain regions and liver. **A-E.** Targeted quantitation heatmaps of BMPs and LPGs from targeted whole-tissue lipidomics of *CLN5*^+/+^, *CLN5*^+/−^, and *CLN5*^−/−^ sheep: primary motor cortex (A), primary visual cortex (B), thalamus (C), cerebellum (D), and liver (E). *CLN5*^+/+^ n = 1, *CLN5*^+/−^ n = 2, *CLN5*^−/−^ n = 3. *CLN5*^+/+^ and *CLN5*^+/−^ have been grouped together for analysis as both animals present as healthy controls. **F.** Targeted quantitation heatmaps of BMPs and LPGs from targeted whole-tissue lipidomics of *CLN5*^+/−^ and *CLN5*^−/−^ sheep: parieto-occipital cortex (F). *CLN5*^+/−^ n = 4, *CLN5*^−/−^ n = 3. Data from targeted QQQ LC/MS. Fold change (FC) by ratio of means. Targeted quantitation heatmaps normalized by PC(18:0_18:1).

### CLN5 Batten disease patient-derived fibroblasts, plasma, and dried blood spots (DBS) have significantly decreased BMP and increased LPG

Given the conserved BMP depletion and LPG accumulation seen across CLN5 Batten disease models, we turned to human patients to assess the utility of BMP and LPG as biomarkers for the disease. We obtained clinical samples from CLN5 Batten disease patients in the form of skin cell-derived fibroblasts, plasma, and DBS, and measured BMP and LPG. Fibroblasts derived from two CLN5 Batten disease patients (c.990G>T (p.Trp330Cys), homozygous; c.227T>G (p.Phe76Cys), homozygous; Table 1) showed a significant decrease in BMP and increase in LPG relative to healthy control fibroblasts (Fig. 5A-D), with those derived from patient #1 showing near complete depletion of BMP. Encouraged by these results, we then analyzed patient samples that would be more clinically useful for biomarker analysis: plasma and DBS. Compared to plasma from healthy individuals, plasma from three CLN5 Batten disease patients (c.448C>T (p.Arg150*), homozygous; c.525del (p.Trp175*), homozygous; c.525del (p.Trp175*), c.641T>A (p.Val214Glu); Table 1) had significantly decreased levels of BMP(18:1_18:1) (Fig. 6A) and concomitantly elevated levels of LPG(18:1) (Fig. 6B). Given the biochemical nature of CLN5 as a BMP synthase utilizing the precursor LPG, a sample-specific calculation of LPG(18:1) peak area divided by BMP(18:1_18:1) peak area led to further increased separation between CLN5 Batten disease patients and controls (Fig. 6C). DBS from three CLN5 Batten disease patients (c.448C>T (p.Arg150*), homozygous; c.525del (p.Trp175*), c.641T>A (p.Val214Glu); c.227T>G (p.Phe76Cys), homozygous; Table 1) also had significantly decreased BMP(18:1_18:1) (Fig. 6D) and significantly increased LPG(18:1) (Fig. 6E). Again, sample-specific LPG/BMP calculation increased the separation between patient samples and controls (21–33-fold) (Fig. 6F). Notably, the BMP depletion observed in DBS samples was more stark than seen in plasma samples, and could be attributed to the inclusion of blood cells in DBS when compared to the extracellular nature of plasma. Overall, these results provide substantial evidence that BMP reduction and LPG elevation are not only phenotypes of cellular, murine, and ovine models, but are also represented in human patients with CLN5 Batten disease. Given the simple collection procedures for plasma and DBS from patients and the robustness of the analytical methods for detecting BMP and LPG, these lipids can serve as important clinical biomarkers for the diagnosis, monitoring, and treatment of CLN5 Batten disease patients.

**Figure 5.**
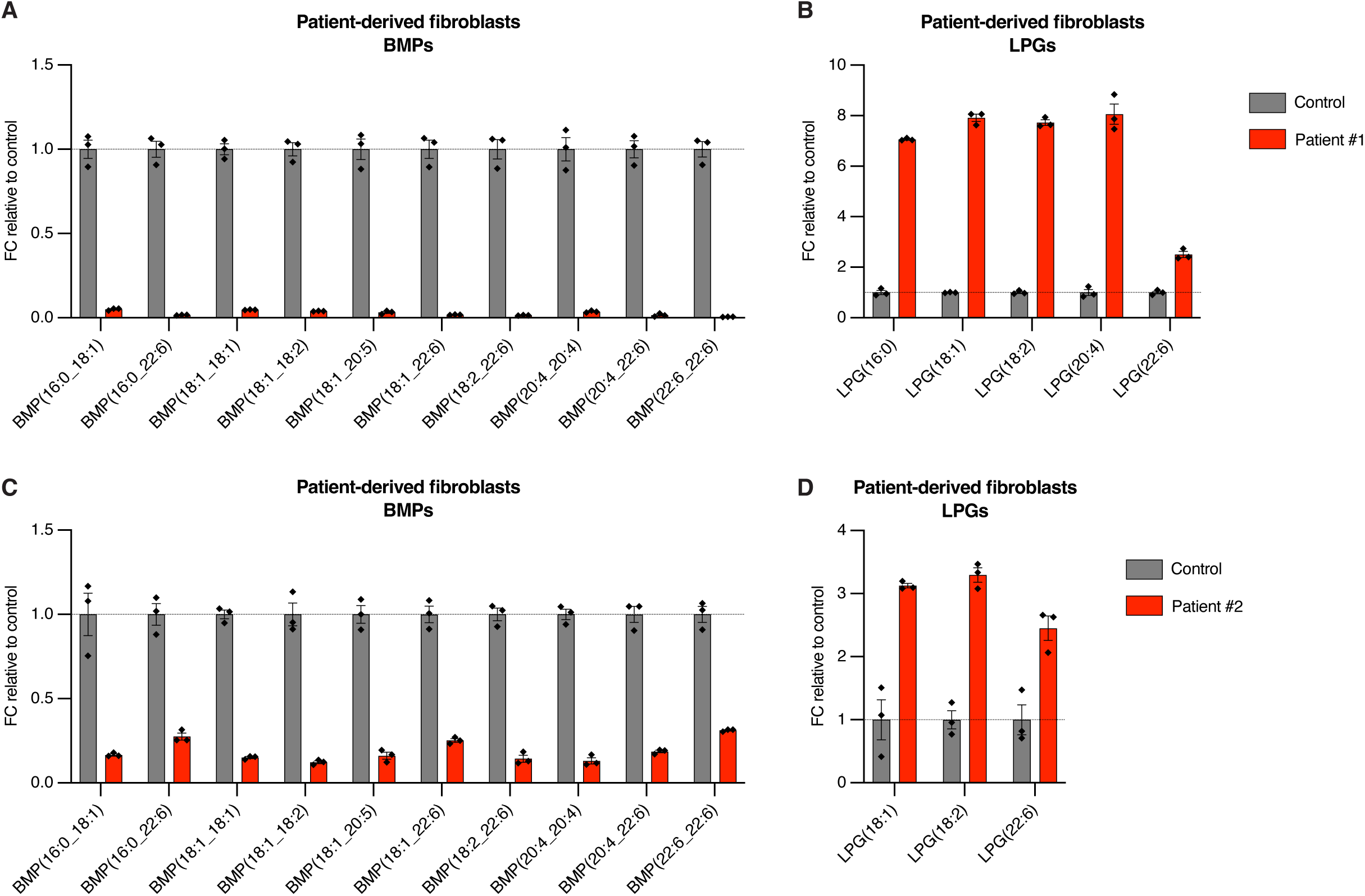
Fibroblasts from patients with CLN5 Batten disease have significantly decreased BMP and increased LPG. **A-B**. Targeted quantitation of biomarker lipids from Hamburg patient #1 fibroblasts compared to control fibroblasts. Control fibroblasts and patient #1 fibroblasts n = 3 technical replicates. **C-D.** Targeted quantitation of biomarker lipids from Hamburg patient #2 fibroblasts compared to control fibroblasts. Control fibroblasts and patient #2 fibroblasts n = 3 technical replicates. Data from targeted QQQ LC/MS. Data presented as fold change (FC) ratio relative to mean of control group. Data plotted as mean ± SEM. Targeted quantitation normalized by PC(18:1_18:1). Patient sample information listed in Table 1.

**Figure 6.**
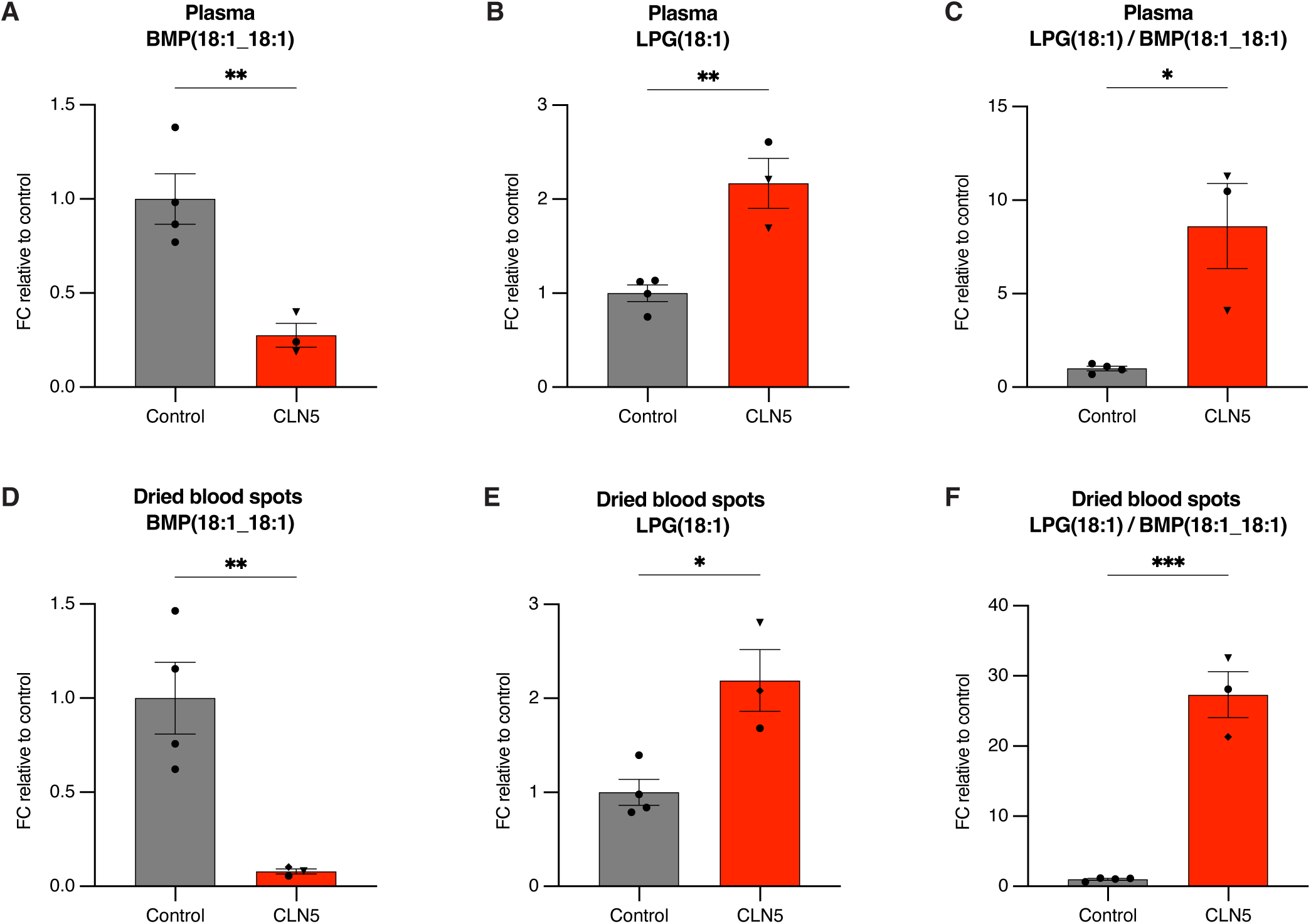
Patients with CLN5 Batten disease have significantly decreased BMP and increased LPG when measured using plasma and dried blood spots. **A-C**. Targeted quantitation of biomarker lipids from patient plasma: BMP(18:1_18:1) (A), LPG(18:1) (B),and ratiometric LPG(18:1)/BMP(18:1_18:1) (C). Healthy control n = 4 (Utrecht n = 4), CLN5 n = 3 (Utrecht n = 2, Pisa n = 1). **D-F.** Targeted quantitation of biomarker lipids from patient dried blood spots (DBS): BMP(18:1_18:1) (A), LPG(18:1) (B), and ratiometric LPG(18:1)/BMP(18:1_18:1) (C). Healthy control n = 4 (Utrecht n = 4), CLN5 n = 3 (Utrecht n = 1, Pisa n = 1, Hamburg n = 1). Data from targeted QQQ LC/MS. Data presented as fold change (FC) ratio relative to mean of control group. Data plotted as mean ± SEM. p-values by Student’s t-tests. Targeted quantitation normalized by PC(18:1_18:1). Statistical comparisons are marked as: * (p < 0.05), ** (p < 0.01), *** (p < 0.001). Patient sample location of collection indicated by symbol shape: Utrecht – circle, Pisa – triangle, Hamburg – rhombus. Patient sample information listed in Table 1.

**Table 1.** Overview of patient information.

## Discussion

The pathological dysregulation of BMP levels in a large number of neurodegenerative diseases underscores the importance of the BMP synthesis pathway to brain and organismal health^33,39–48,52–54^. As such, BMP is becoming increasingly recognized for its potential as a clinical biomarker and therapeutic target^33,49–51,55,56,87^. Insight into the final step of BMP synthesis—namely, its generation from LPG through CLN5-mediated deacylation and transacylation reactions^32^—strongly alludes toward inversely dysregulated levels of BMP and LPG as a clinical biomarker for CLN5 Batten disease. Our results here confirm that loss of functional CLN5 is sufficient to cause significantly decreased BMP and increased LPG abundance in murine and ovine models of CLN5 Batten disease as well as in patient samples. These changes are observable across several distinct biological and clinical materials. In making this assessment, we thoughtfully and rigorously consider the challenges of quantifying BMP in the presence of its structural isomer PG^86^, denoting three distinct confidence levels for annotation of each BMP species. In ovine models of CLN5 Batten disease, lipidomics conducted on multiple brain regions revealed significant reduction of BMP and accumulation of LPG as a feature conserved in mammals, in a region-invariant manner, and among some of the most susceptible regions to neurodegeneration in advanced stages. Studies in these models are of great importance as they represent preclinical models for studying human Batten disease-relevant pathology and evaluating gene therapy^23–31^. Directly showcasing the translatable relevance of BMP and LPG as biomarkers for CLN5 Batten disease, lipidomics from patient-derived fibroblasts showed significant reduction of BMP species and accumulation of LPG species. Finally, to apply these insights to the clinical setting, lipidomics from plasma and dried blood spots both demonstrated significant BMP reduction and LPG accumulation. In conclusion, we propose the combined observance of BMP reduction and LPG accumulation from patient dried blood spots and/or plasma as an accessible biomarker for the diagnosis and monitoring of CLN5 Batten disease^49,55,56,67,79,81^. As treatment options emerge for CLN5 Batten disease, we hope these biomarkers will prove useful for clinicians to effectively and quickly diagnose patients as well as to monitor patient responses to treatment. This work fits within a key translational pipeline linking biochemical functions with biologically– and clinically-relevant contexts by moving from biochemical insights to model organisms and, ultimately, to human patients. As research into the biochemical functions of other proteins encoded by *CLN* genes progresses, this approach is poised to elucidate additional clinically relevant biomarkers to facilitate the treatment of patients with rare genetic diseases.

## Methods

### Sex as a biological variable

Sex was not considered as a biological variable due to the overall small number of mice, sheep, and patients available, limiting grouping by sex. Sexes of all patients are reported (Table 1). The selection and enrollment of patients was mainly based on the feasibility and availability of patients with CLN5 Batten disease, as the condition is very rare.

### Mouse studies

Mice were acquired and maintained as previously described^39,87,88^. Mice were maintained on a standard light–dark cycle with *ad libitum* access to food and water in a room with a controlled temperature (22 °C) and humidity (∼50%). Cages were cleaned every 4–5 days, and water and food supplies were checked daily. All of the procedures involving mice were carried out in accordance with the approved guidelines of the Institutional Animal Care and Use Committee at Stanford University. The LysoTag mouse strain was previously reported (Jackson Laboratory (JAX); strain no. 035401)^39^. The *Cln5*^−/−^ mouse was previously generated^13^. *Cln5*^−/−^ and LysoTag mice were bred to generate a LysoTag-harboring murine model of Batten disease as previously described^39^. Mouse genotyping for the indicated alleles was performed as reported in the original publications.

### Whole-tissue and LysoIP collection of mouse tissues

Lysosome immunoprecipitation (LysoIP) was conducted following established protocols, with adaptation as necessary^39,86–92^. Whole-tissue collection was adapted from the whole-cell/whole-tissue sections of the equivalent LysoIP protocols. For all mouse tissue collections, the tissues of mice were obtained after euthanasia and used immediately without freezing for whole-tissue extraction or LysoIP.

For whole-tissue collection for lipidomics, uniform tissue samples collected from each mouse were: half brain, whole heart, one kidney, 4-mm-diameter liver biopsy punch, and one lung. Each tissue sample was weighed, to ensure relative mass uniformity, and then homogenized (DWK Life Sciences, cat. no. K749540-0000) on ice with 50 μL of potassium phosphate buffered saline (KPBS; 136 mM KCl, 10 mM KH_2_PO_4_, pH = 7.25 in Optima LC-MS water). 50 μL of the homogenized slurry was then transferred to a new tube for lipid extraction, at which point 1 mL of lipid extraction buffer was added (for downstream lipid extraction, see “Lipid extraction from all sample types”).

For whole-tissue collection for enzyme assays, the uniform tissue sample collected from each mouse was a 4-mm-diameter liver biopsy punch each. Each tissue sample was weighed, to ensure relative mass uniformity, and then dounced (VWR, cat. no. 89026-386) on ice with 1 mL of 20 mM HEPES (Gibco, cat. no. 15630080) buffer (pH = 7.4) with 150 mM NaCl and 0.05% LMNG (Anatrace, cat. no. NG310 5 GM). 400 μL of the homogenized slurry was then transferred to a new tube, immediately snap frozen with liquid nitrogen, and stored at −80 °C for later enzyme assays (for downstream assay, see “Lysosomal lysate and whole-tissue homogenate enzyme assays”).

For LysoIP collection for lipidomics, the uniform tissue samples collected from each mouse were half brain and 4-mm-diameter liver biopsy punch. Each tissue sample was weighed, to ensure relative mass uniformity, and then dounced on ice with 1 mL of KPBS. The remaining downstream steps were directly adapted from previous LysoIP protocols. In brief, the 2 mL slurry was centrifuged at 1,000*g* for 2 min at 4 °C. Then, 1 mL of the supernatant was incubated with 200 μL anti-HA magnetic beads (Thermo Scientific 88837) on a laboratory rocker for 15 min at 4 °C, and subsequently washed three times with 1 mL KPBS per wash using a DynaMag Spin Magnet (Invitrogen, cat. no. 12320D) to aspirate liquid after each wash. Finally, 1 mL of lipid extraction buffer was added to the isolated lysosome-bound beads (for downstream lipid extraction, see “Lipid extraction from all sample types”).

For LysoIP collection for enzyme assays, the uniform tissue samples collected from each mouse were full brain and one large liver lobe. Each sample was weighed, to ensure relative mass uniformity. Brain samples were then dounced on ice with 1 mL of phosphate buffered saline (PBS; Gibco, cat. no. 10010031) with cOmplete protease inhibitor cocktail (Roche, cat. no. 11697498001) in standard size douncers (VWR, cat. no. 89026-386). After douncing, another 1 mL of PBS with protease inhibitor cocktail (PBSi) was added to the mixture prior to centrifugation. Liver samples were dounced on ice with 10 mL of PBSi in large size douncers (VWR, cat. no. 89026-382). The remaining downstream steps were directly adapted from previous LysoIP protocols. In brief, the 2 mL brain and 10 mL liver slurries were centrifuged at 1,000*g* for 2 min at 4 °C, the supernatants were incubated with 200 μL anti-HA magnetic beads (Thermo Scientific 88837) for brain and 1 mL anti-HA beads magnetic beads for liver on a rotator shaker for 15 min at 4 °C, and were all subsequently washed three times with 1 mL PBSi per wash using a DynaMag Spin Magnet (Invitrogen, cat. no. 12320D) to aspirate liquid after each wash. Finally, 50 μL of ultrapure distilled water was added to the isolated lysosome-bound beads and the mixture was incubated for 30 min at 37 °C. Using a DynaMag Spin Magnet, the supernatant was then transferred to a new tube, immediately snap frozen with liquid nitrogen, and stored at −80 °C for later enzyme assays (for downstream assay, see “Lysosomal lysate and whole-tissue homogenate enzyme assays”).

### Whole-tissue collection of sheep tissues

Tissue samples from sheep were obtained in Lincoln, New Zealand as previously described^28,93^. In brief, multiple brain regions and liver were obtained from sheep within 5 minutes after euthanasia. Samples were immediately snap frozen without further processing and stored at −80 °C. The samples were then shipped to Stanford, CA on dry ice, where they were received and confirmed to still be on dry ice, removed, and stored at −80 °C. On the day of extraction, samples were thawed on ice and then aliquoted using knife and scissors. Specifically, small uniform sections of each tissue were cut and weighed to ensure relative mass uniformity of approximately 75 mg for primary motor cortex, primary visual cortex, thalamus, cerebellum, and liver and approximately 35 mg for parieto-occipital cortex. Each tissue sample was then dounced on ice with 1 mL of KPBS. 200 μL of the homogenized slurry was then transferred to a new tube for lipid extraction, at which point 1 mL of lipid extraction buffer was added (for downstream lipid extraction, see “Lipid extraction from all sample types”).

### Fibroblasts, plasma, and dried blood spots collection from patients

Fibroblasts were derived from two CLN5 Batten disease patients in Hamburg, Germany as previously described^94–96^. Control and patient fibroblasts were cultured in Dulbecco’s Modified Eagle Medium (DMEM; Gibco, cat. no. 11965118) with 10% heat inactivated fetal bovine serum (FBS; Gibco, cat. no. 10438026) and 1% penicillin-streptomycin (Cytiva, cat. no. SV30010). For lipid extraction, fibroblasts were collected from culture as three cell pellets of 1×10^6^ cells each by trypsinization, cell counting, and pelleting. 1 mL of lipid extraction buffer was then added to each technical replicate cell pellet (for downstream lipid extraction, see “Lipid extraction from all sample types”).

Plasma was collected from patients in Utrecht, The Netherlands and Pisa, Italy as previously described^97,98^. Plasma samples were shipped on dry ice to Stanford, CA, where they were stored at −80 °C. On the day of extraction, samples were thawed on ice. 50 μL of plasma from each sample was transferred to a new tube and 1 mL of lipid extraction buffer was added to each plasma sample aliquot (for downstream lipid extraction, see “Lipid extraction from all sample types”).

Dried blood spots (DBS) were collected from patients in Utrecht, The Netherlands; Pisa, Italy; and Hamburg, Germany as previously described^98,99^. Samples were shipped on dry ice to Stanford, CA, where they were stored at −80 °C. On the day of extraction, samples were thawed on ice. Lipid extraction from DBS was adapted from previously described methods^100,101^. A circular cut out of a single dried blood spot was cut out using scissors for each patient and placed in a new tube. 1 mL of lipid extraction buffer was then added to each tube containing a dried blood spot cut out (for downstream lipid extraction, see “Lipid extraction from all sample types”).

### Lysosomal lysate and whole-tissue homogenate enzyme assays

Enzyme assays were conducted using an adapted method^32,87^. On an intermediate day prior to enzyme assays, all lysosomal lysate and whole-tissue homogenate samples were measured for protein concentration by bicinchoninic acid (BCA) assay. Samples (previously collected, snap frozen, and stored at −80 °C) were quickly thawed at 37 °C and placed on ice. Using a Pierce BCA Protein Assay Kit (Thermo Scientific, cat. no. 23225), Pierce BCA Reagent A was mixed with Pierce BCA Reagent B in a 50:1 (v/v) ratio. 200 μL of the mixed working reagent was then placed in each well of a 96-well plate. 2 μL of lysosomal lysate and whole-tissue homogenate samples were added to one well each. 2 μL of premade serial dilutions of bovine serum albumin in appropriate buffer were also added to one well each as a standard curve. The appropriate buffer for lysosomal lysate measurement was ultrapure distilled water. The appropriate buffer for whole-tissue homogenate measurement was 20 mM HEPES (Gibco, cat. no. 15630080) buffer (pH = 7.4) with 150 mM NaCl and 0.05% LMNG (Anatrace, cat. no. NG310 5 GM). The standard curve concentrations of bovine serum albumin were: 0, 25, 125, 250, 500, 750, 1000, 1500, and 2000 μg/mL. The 96-well plate was then incubated in aluminum foil at 37 °C for 30 min, after which it was read on an absorbance microplate reader according to manufacturer instructions. Protein concentrations were then calculated for all lysosomal lysate and whole-tissue homogenate samples.

All lysosomal lysate and whole-tissue homogenate enzyme assays were performed as 100 μL reactions with 25 μM d5-LPG(17:0) substrate (Avanti, cat. no. 858130L-1mg). On the day of enzyme assays, 100 μL of 100 μM pre-diluted stock d5-LPG(17:0) was placed into reaction tubes and dried in a SpeedVac vacuum concentrator (Labconco, cat. no. 7315060) at room temperature. 80 μL reaction buffer was then added to reaction tubes and the LPG substrate was resuspended by sonicating in a sonicator bath for 1 minute followed by centrifuging in a tabletop mini centrifuge for 30 seconds. For acidic pH reactions, the reaction buffer was 50 mM sodium acetate-acetic acid (pH = 5.5) with 150 mM sodium chloride. For neutral pH reactions, the reaction buffer was phosphate buffered saline (PBS) (pH = 7.4). Resuspended reaction tubes were then placed on ice. Lysosomal lysate or whole-tissue homogenate samples (previously collected, snap frozen, and stored) were then quickly thawed at 37 °C and placed on ice. Using the precalculated protein concentrations, lysosomal lysate and whole-tissue homogenate samples were added such that the final protein mass in each tube was 5 μg. For rCLN5 rescue experiments, reactions were also supplemented with recombinant human CLN5 protein (rCLN5)^32^ such that the final rCLN5 concentrations were 5 nM or 10 nM. Each reaction was then topped off with its pH-respective buffer such that the final reaction volume in each tube was 100 μL. All enzyme assay experiments had negative controls of buffer alone, buffer with LPG substrate and without biological material, and buffer with biological material and without LPG substrate; all volumes were adjusted accordingly. Following reaction set up, a 20 μL aliquot of each reaction was transferred into a pre-prepared tube containing 1 mL lipid extraction buffer (LEB) as the 0 min time point of said reaction. All reaction tubes were then moved from ice to 37 °C and reaction time began. Analogous 20 μL aliquots were transferred to 1 mL LEB tubes at the appropriate reaction time points, which included 30 min, 1 h, 2 h, and 4 h. All 20 μL aliquots in 1 mL LEB underwent lipid extraction (for downstream lipid extraction, see “Lipid extraction from all sample types”).

For lysosomal lysate enzyme assays, BMP synthesis was monitored using targeted QQQ LC/MS measurement of synthesized d5-BMP(17:0_17:0). For whole-tissue homogenate enzyme assays, BMP synthesis was monitored using targeted QQQ LC/MS measurement of synthesized d5-BMP(17:0_18:1), which was found to be more abundant in whole-tissue experiments. Peak area was normalized using endogenous PC(18:1_18:1) (DOPC) with monitoring of substrate d5-LPG(17:0), internal standard d7-PG(15:0_18:1), and endogenous BMPs, LPGs, PGs, and PCs for quality checks.

### Lipid extraction from all sample types

Lipids were extracted using a previously described method^86,91,102,103^, with adaptation as necessary for different sample types. Lipids were harvested from various biological samples with 1 mL lipid extraction buffer (LEB), 2:1 chloroform:methanol (v/v) supplemented with 750 ng/mL SPLASH LIPIDOMIX isotopically labeled internal standards (Avanti, cat. no. 330707-1EA), by vortexing at 4 °C for 1 h, and further extracted after the addition of 200 μL 0.9% NaCl saline solution (Teknova, cat. no. S5819) by vortexing at 4 °C for 10 min. The mixture was then centrifuged at 3,000*g* and 4 °C for 5 minutes, after which 600 μL of the bottom chloroform phase was collected, dried in a SpeedVac vacuum concentrator (Labconco, cat. no. 7315060) at room temperature, and stored at −80°C. Samples were reconstituted in 50 μL 13:6:1 acetonitrile:isopropanol:water (v/v/v) solution by vortexing at 4 °C for 10 min and then centrifuged at 30,000*g* and 4 °C for 15 minutes. The supernatant was then transferred to glass LC/MS vials for analysis by LC/MS lipidomics.

For whole-tissue collection of mouse tissue, lipids were extracted from 50 μL homogenized whole-tissue slurry with 1 mL LEB. For whole-tissue collection of sheep tissue, lipids were extracted from 200 μL homogenized whole-tissue slurry with 1 mL LEB. For LysoIP collection of mouse tissue, lipids were extracted from isolated lysosome-bound beads with 1 mL LEB. As an added step for LysoIP beads, the mixture containing beads and 1 mL LEB was placed on a DynaMag Spin Magnet halfway through its 1 h vortexing extraction, and the supernatant was transferred to a new tube. Vortexing extraction of the supernatant was then continued for the remaining 30 min, and proceeded through the remaining lipid extraction procedure otherwise unmodified. For patient fibroblasts, lipids were extracted from 1×10^6^ cell pellets with 1 mL LEB.

For patient plasma, lipids were extracted from 50 μL of plasma with 1 mL LEB. For patient DBS, lipids were extracted from circular cut outs of one DBS each with 1 mL LEB. At the end of the first 1 h vortexing extraction step, the circular DBS cut out was manually removed from each tube using metal forceps. The extraction procedure then proceeded otherwise unmodified, including the subsequent saline addition step. For lysosomal lysate and whole-tissue homogenate enzyme assays, lipids were extracted from 20 μL reaction aliquots with 1 mL LEB.

### Untargeted LC/MS lipidomics

Untargeted LC/MS lipidomics was performed following a previously described protocol^86^. Liquid chromatography was performed using a Vanquish HPLC (Thermo Fisher Scientific) with an Ascentis Express C18 150 × 2.1 mm column (Millipore Sigma, cat. no. 53825-U) coupled with a 5 × 2.1 mm guard assembly (Sigma-Aldrich, cat. no. 53500-U, cat. no. 50542-U). Mobile phase A was 60% water and 40% acetonitrile (v/v) containing 10 mM ammonium formate and 0.1% formic acid (v/v). Mobile phase B was 90% isopropanol and 10% acetonitrile (v/v) containing 10 mM ammonium formate and 0.1% formic acid (v/v). The flow rate was 0.26 mL/min. The elution gradient was as follows: 32% B 0–1.5 min, 45% B 1.5–4 min, 52%B 4–5 min, 58% B 5–8 min, 66% B 8–11 min, 70% B 11–14 min, 75% B 14–18 min, 97% B 18–21 min, 97% B 21–35 min, 32% B 35–35.1 min, 32% B 35.1–40 min. Mass spectrometry was performed using an Orbitrap ID-X Tribrid mass spectrometer (Thermo Fisher Scientific) with heated electrospray ionization. The Orbitrap was operated separately on both positive and negative mode, with each sample injected twice, once for each polarity. Data was acquired using both MS1 and data-dependent MS2 scanning.

### Untargeted LC/MS lipidomics analysis

Untargeted Orbitrap LC/MS lipidomics data was analyzed using a previously described protocol^86^. For the generation of untargeted volcano plots, lipids were first annotated by LipidSearch 4.2 (LS) (Thermo Fisher Scientific) using MS2 data. Lipids were then quantified by Compound Discoverer 3.3 (CD) (Thermo Fisher Scientific) using MS1 data and annotations by LS. All peak areas were normalized by CD using constant median normalization. Peak areas were then filtered to only include peaks with lipid annotation names by LS, |annotation delta mass| ≤ 5 ppm, and signal-to-noise ratio ≥ 3. Filtered lipids were then plotted in a volcano plot using fold change (FC) and p-values by Student’s t-tests. For the generation of targeted quantitation heatmaps, MS2-dependent lipid annotations from LS were used to manually identify lipids in FreeStyle 1.5 (FS) (Thermo Fisher Scientific) and quantify them in TraceFinder (TF) (Thermo Fisher Scientific). BMPs, PGs, LPGs, and PEs were manually quantified using TF as [M−H]^−^ ions on negative polarity with 5 ppm mass tolerance. All target lipids and normalizers were filtered for signal-to-noise ratio ≥ 3 and pooled quality control linearity check R^2^ ≥ 0.90. BMPs, PGs, and LPGs were then normalized using endogenous PE(18:1_18:1) (DOPE) and displayed in a heatmap using FC. Volcano plot highlighted lipids, including all BMPs, PGs, and LPGs, and all heatmap lipids were manually and rigorously validated for proper annotation by examining MS1 and MS2 spectra in LS, CD, and FS. All BMPs, PGs, and LPGs were ensured to be quantified as [M−H]^−^ in CD, and to have proper MS1 peak integration visually. All LPGs were ensured to have proper MS2 spectra and retention time (RT) relative to other LPG acyl chains, ensuring reliable annotation. All PEs were ensured to have proper MS2 spectra and retention time (RT) relative to other PE acyl chains, ensuring reliable annotation. All BMPs and PGs were examined for [M−H]^−^ and [M+NH_4_]^+^ MS2. [M−H]^−^ MS2 were used to ensure reliable annotation as PG/BMP, non-distinguishing. [M+NH_4_]^+^ MS2 were used to ensure reliable annotation as either BMP or PG, distinguishing, based on [M+NH_4_]^+^ MS2 fragmentation to MG or DG, respectively. All BMPs and PGs were also ensured to have proper RT relative to other BMP and PG acyl chains as well as isomeric BMP and PG. In each tissue, BMPs and PGs with both [M−H]^−^ and [M+NH_4_]^+^ MS2 annotation in the tissue of interest were labeled as confidence level 1 (cl1). BMPs and PGs with [M−H]^−^ MS2 annotation in the tissue of interest, and [M+NH_4_]^+^

MS2 annotation of an isomeric BMP or PG in that same tissue, were labeled as confidence level 2 (cl2). In the untargeted volcano plots, all other BMPs and PGs that had only [M−H]^−^ MS2 annotation in the tissue of interest were labeled as PG/BMP. In the targeted quantitation heatmaps, BMPs and PGs with [M−H]^−^ MS2 annotation in the tissue of interest, and [M+NH_4_]^+^ MS2 annotation of the same or an isomeric BMP or PG in a different tissue, were labeled as confidence level 3 (cl3). In the targeted quantitation heatmaps, BMPs and PGs without cl1, cl2, or cl3 were not considered to have minimally confident annotation, and were not quantified.

### Targeted LC/MS lipidomics

Targeted LC/MS lipidomics was performed following a previously described protocol^86^. Liquid chromatography was performed using a 1290 Infinity II HPLC (Agilent Technologies) with an Agilent RRHD Eclipse Plus C18, 2.1 mm, 100 mm, 1.8 µm column coupled with a C18, 2.1 mm, 1.8 µm guard (Agilent Technologies, cat. no. 821725-901. Mobile phase A was 60% water and 40% acetonitrile (v/v) containing 10 mM ammonium formate and 0.1% formic acid (v/v). Mobile phase B was 90% isopropanol and 10% acetonitrile (v/v) containing 10 mM ammonium formate and 0.1% formic acid (v/v). The flow rate was 0.4 mL/min. The elution gradient was as follows: 15% B 0–3 min, 70% B 3–14 min, 75% B 14–15 min, 100% B 15–15.2 min, 15% B 15.2–16 min, followed by 15% B posttime for 2 min. Mass spectrometry was performed using an Ultivo QQQ mass spectrometer (Agilent Technologies) with AJS-electrospray ionization. Lipids were measured in a targeted manner using multiple reaction monitoring (MRM). MRM transitions were specifically designed for each lipid class. MRM transitions for BMP were from the positive ammonium adduct precursor ions ([M+NH_4_]^+^) to BMP-specific monoacylglycerol (MG) fragments ([MG−H_2_O+H]^+^). MRM transitions for PG were from the positive ammonium adduct precursor ions ([M+NH_4_]^+^) to PG-specific diacylglycerol (DG) fragments ([DG−H_2_O+H]^+^). MRM transitions for LPG were from the negative deprotonated parent precursor ion ([M−H]^−^) to fatty acid fragments ([FA−H]^−^) and glycerophosphate fragments ([GP−H_2_O−H]^−^). MRM transitions for PC were from the positive protonated parent precursor ion ([M+H]^+^) to lysophosphatidylcholine fragments ([LPC−H_2_O+H]^+^), phosphocholine fragments ([PCho]^+^), and choline fragments ([Cho−H_2_O]^+^). Deuterium labeling for enzyme assays as well as isotopically labeled internal standards was additionally accounted for in MRM transitions where necessary.

### Targeted LC/MS lipidomics analysis

Targeted QQQ LC/MS lipidomics data was analyzed using a previously described protocol^86^. Lipids were first examined for annotation using MassHunter Qualitative Analysis (Agilent Technologies). All lipids were checked for proper RT relative to past and other present data, as well as to frequently run lab lipid standards. All lipids were also checked for proper RT relative to other lipids within the same data set. Lipids with multiple MRM transitions were checked for reliability across multiple transitions. Lipids, including BMPs, PGs, LPGs, and PCs, were then manually quantified using MassHunter QQQ Quantitative Analysis (Quant-My-Way) (Agilent Technologies). BMPs were quantified with MRM transition [M+NH_4_]^+^ to [MG−H_2_O+H]^+^, and qualified with an alternate MG transition where applicable. PGs were measured with MRM transition [M+NH_4_]^+^ to [DG−H_2_O+H]^+^. LPGs were measured with MRM transition [M−H]^−^ to [FA−H]^−^, and qualified with transition to [GP−H_2_O−H]^−^. PCs were measured with MRM transition [M+H]+ to [PCho]^+^, and qualified with transitions to [Cho−H_2_O]^+^, [LPC−H_2_O+H]^+^, and an alternate LPC where applicable. All target lipids and normalizers were filtered for signal-to-noise ratio ≥ 3 and pooled quality control linearity check R^2^ ≥ 0.90. BMPs, PGs, and LPGs were then normalized using endogenous PC(18:1_18:1) (DOPC), or endogenous PC(18:0_18:1) in cases where DOPC did not have linearity check R^2^ ≥ 0.90. Lipids were then displayed in xy plots as normalized peak area and heatmaps and bar graphs using FC.

### Statistics

Lipid Search 4.2 (Thermo Fisher Scientific), Compound Discoverer 3.3 (Thermo Fisher Scientific), FreeStyle 1.5 (Thermo Fisher Scientific), and TraceFinder 5.1 (Thermo Fisher Scientific) were used for untargeted Orbitrap LC/MS lipidomics raw data analysis. MassHunter Qualitative Analysis (Agilent Technologies) and MassHunter QQQ Quantitative Analysis (Quant-My-Way) (Agilent Technologies) were used for targeted QQQ LC/MS lipidomics raw data analysis. Exported numerical data was further processed and analyzed using Microsoft Excel 365 and Prism 10 (GraphPad Software). Data was visualized using Prism 10 and Adobe Illustrator. Figure 3A was created using BioRender (https://www.biorender.com/). The statistical test used throughout for comparison between two groups was an independent samples t-test with pooled variance (Student’s t-test, unpaired, homoscedastic). Statistical tests were only performed on groups with biological replicates, and not performed on groups with technical replicates. All bar graph data are expressed as mean ± standard error of the mean (SEM). All p-values < 0.05 were considered significant. Where applicable, p-values were represented by asterisks: ns (not significant), * (p < 0.05), ** (p < 0.01), *** (p < 0.001), **** (p < 0.0001).

### Study approval

Patient participation from Hamburg, Germany and study was approved by the respective local medical ethics committee of the Ärztekammer Hamburg, Germany (PV7215). Patients from Pisa, Italy were recruited through the Italian NCL Registry (ClinicalTrials.gov identifier NCT06844877). The protocol was approved by the Tuscany Regional Pediatric Ethics Committee (CEPR; approval no. 130/2024; amendment no. 198/2024). Patients from Utrecht, the Netherlands consented to use their residual material collected for diagnostic purposes in the Wilhelmina Children’s Hospital metabolic biobank (TCBio 19–489/B https://tcbio.umcutrecht.nl).

### Data availability

All data needed to evaluate the conclusions in the paper are present in the manuscript and supplemental material. Supporting data values associated with the main article and supplemental material are included in the Supporting Data Values file, with separate tabs for each applicable figure panel. All LC/MS raw data will be archived in general purpose repositories with free access following peer review and upon publication.

## Author contributions

The study was conceived by ESR, NM, and MA-R. Methodology was developed by ESR, NM, HNA, and MA-R. Experiments were conducted by ESR, NM, HNA, MI, IVP, CP, and LMW. Sheep samples were acquired by SJM and NLM. Patient samples were acquired by CP, LMW, DMP, SDV, SD, MM, FMS, PMvH, and AS. Data acquisition was conducted by ESR, NM, HNA, and WD. Data analysis, data visualization, and figure harmonization were performed by ESR. Funding was acquired by FMS, PMvH, NLM, AS, and MA-R. Project administration and supervision were conducted by MA-R. The original draft was written by ESR, NM, and MA-R. Reviewing and editing was done by all authors. Author order was determined by mutual agreement.

## Acknowledgements

We thank all the members of the Abu-Remaileh Laboratory for their helpful discussions and the Metabolomics group (directed by Dr. Yuqin Dai) of the Nucleus at Sarafan ChEM-H, Stanford University. We also thank Esther Sammler for discussion and advice, and Katja Kanninen from The University of Eastern Finland for sharing the *Cln5*^−/−^ mouse model.

## Supplementary material

**Figure S1.** *Cln5*^−/−^ mice exhibit minimal change of PGs across tissues and in lysosomes. **A-E**. Targeted quantitation heatmaps of PGs from untargeted whole-tissue lipidomics, corresponding to Fig. 1A-E volcano plots: brain (A), heart (B), kidney (C), liver (D), and lung (E). *Cln5*^−/−^ mice n = 5, *Cln5*^+/−^ mice n = 10 for A-D, *Cln5*^+/−^ mice n = 9 for E. **F-G.** Targeted quantitation heatmaps of PGs from untargeted LysoIP lipidomics, corresponding to Fig. 2A-B volcano plots: brain (F) and liver (G). *Cln5*^+/−^ mice n = 4, *Cln5*^−/−^ mice n = 3. Data from untargeted Orbitrap LC/MS. Fold change (FC) by ratio of means. p-values by Student’s t-tests. Targeted quantitation heatmaps normalized by PE(18:1_18:1). All quantified species in heatmaps were manually and rigorously validated for proper annotation and integration. PGs with confidence level 1 (cl1) label are annotated by definitive +NH_4_ MS2 and −H MS2. PGs with cl2 label are annotated by −H MS2 and relative retention time (RT) to an isomeric cl1 BMP in the same tissue. PGs with cl3 label are annotated by −H MS2 and relative RT to an isomeric cl1 BMP in a different tissue.

**Figure S2.** *Cln5*^+/+^ and *Cln5*^+/−^ mice exhibit no difference in BMPs, LPGs, and PGs. **A-O**. Targeted quantitation heatmaps of BMPs, LPGs, and PGs from targeted whole-tissue lipidomics of *Cln5*^+/+^ and *Cln5*^+/−^ mice: brain (A-C), heart (D-F), kidney (G-I), liver (J-L), and lung (M-O). *Cln5*^+/+^ mice n = 3, *Cln5*^+/−^ mice n = 14 for A-F and J-O, *Cln5*^+/−^ mice n = 13 for G-I. Data from targeted QQQ LC/MS. Fold change (FC) by ratio of means. Targeted quantitation heatmaps normalized by PC(18:1_18:1). p-values by Student’s t-tests. Targeted quantitation normalized by PC(18:1_18:1). Statistical comparisons are marked as: ns (not significant), * (p < 0.05).

**Figure S3.** *CLN5*^−/−^ sheep exhibit minimal change of PGs across brain regions and liver. **A-E**. Targeted quantitation heatmaps of BMPs and LPGs from targeted whole-tissue lipidomics of *CLN5*^+/+^, *CLN5*^+/−^, and *CLN5*^−/−^ sheep, corresponding to Fig. 4A-E heatmaps: primary motor cortex (A), primary visual cortex (B), thalamus (C), cerebellum (D), and liver (E). CLN5^+/+^ n = 1, *CLN5*^+/−^ n = 2, *CLN5*^−/−^ n = 3. *CLN5*^+/+^ and *CLN5*^+/−^ have been grouped together for analysis as both animals present as healthy controls. F. Targeted quantitation heatmaps of BMPs and LPGs from targeted whole-tissue lipidomics of *CLN5*^+/−^ and *CLN5*^−/−^ sheep, corresponding to Fig. 4F heatmap: parieto-occipital cortex (F). *CLN5*^+/−^ n = 4, *CLN5*^−/−^ n = 3. Data from targeted QQQ LC/MS. Fold change (FC) by ratio of means. Targeted quantitation heatmaps normalized by PC(18:0_18:1).

**Table S1.** Untargeted lipidomics of *Cln5*^+/−^ and *Cln5*^−/−^ mice tissues.

