## Supplementary figures and images for "Translational lipidomics reveals BMP and its precursor LPG as biomarkers for CLN5 Batten disease"

### Supplementary Figure 1

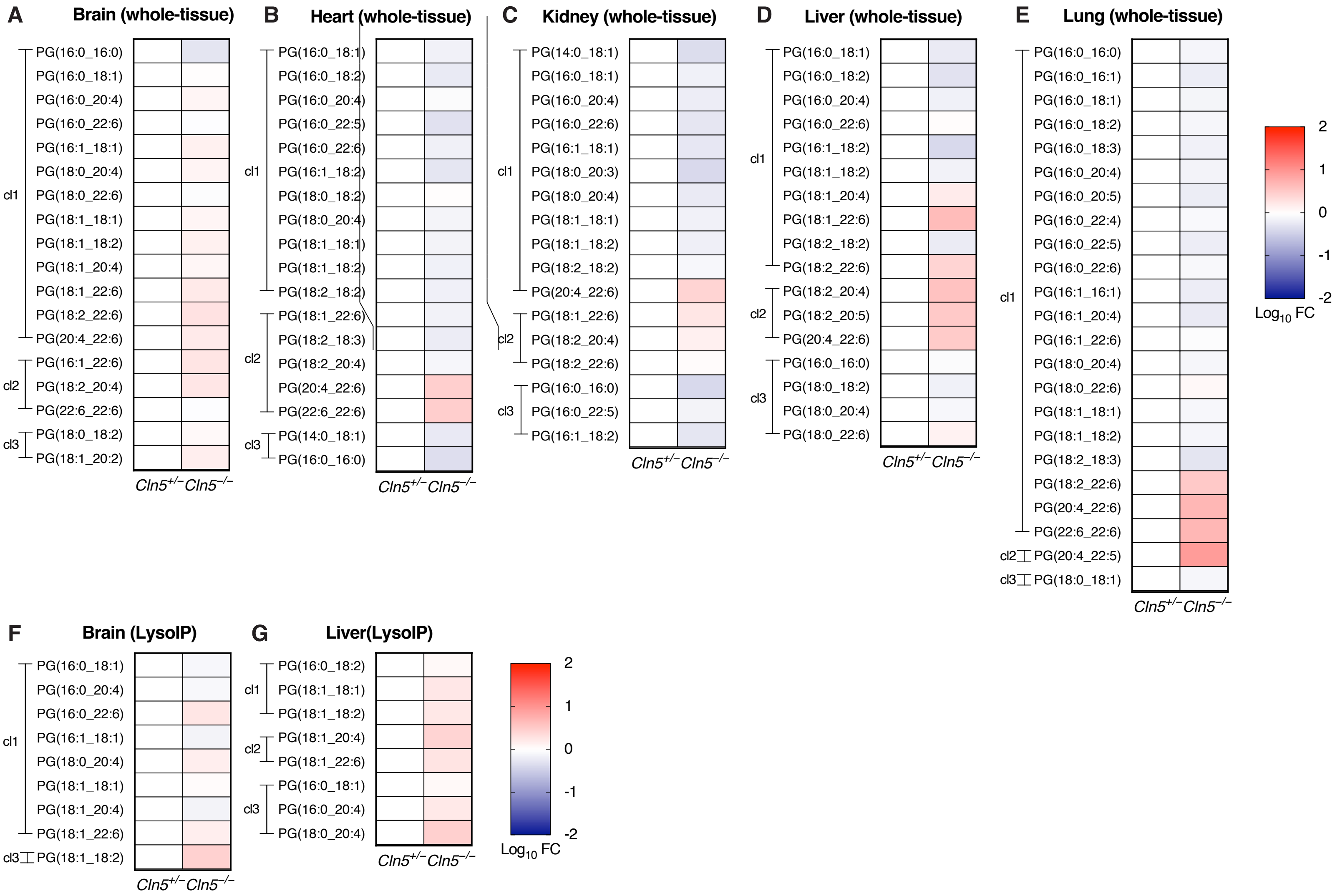

### Supplementary Figure 2

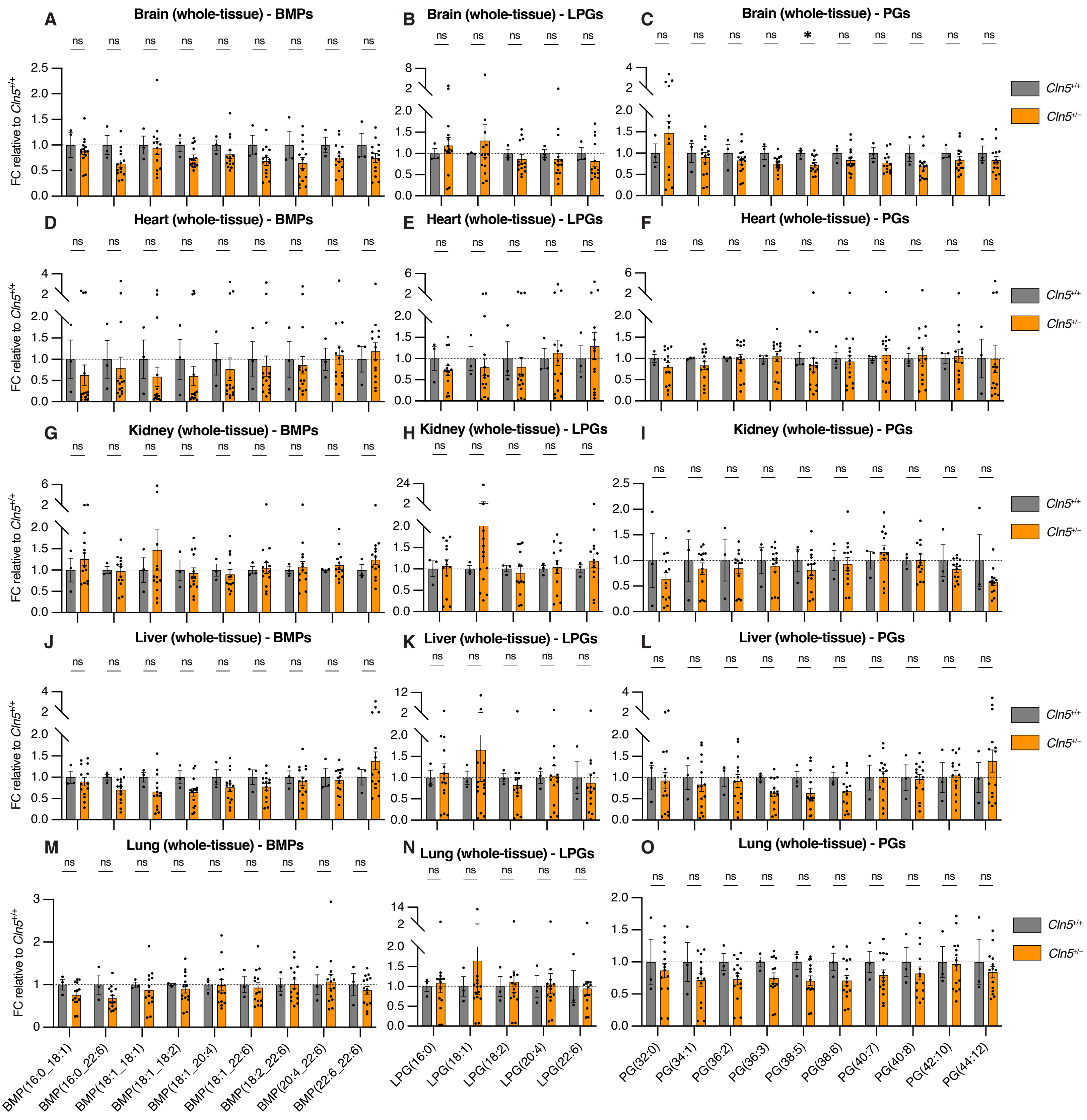

### Supplementary Figure 3

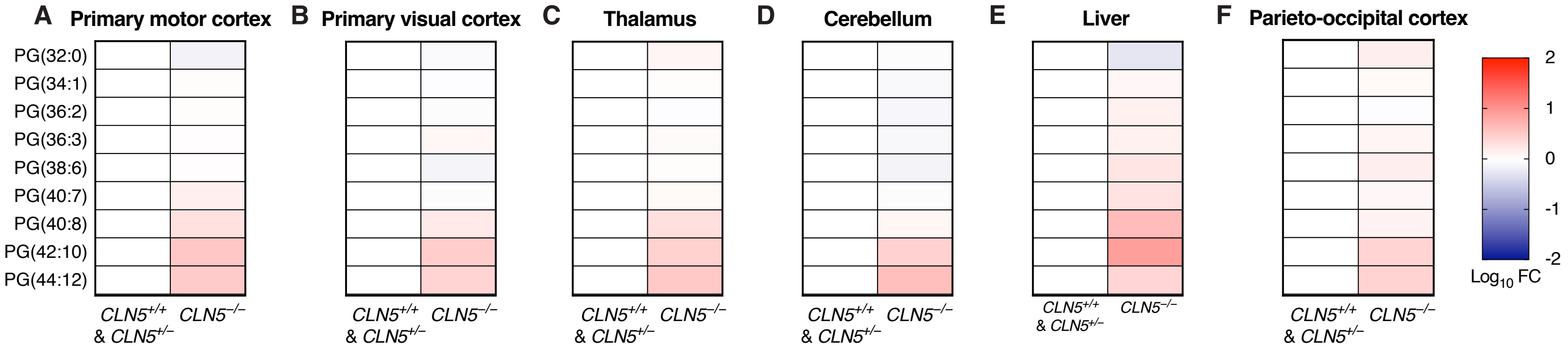
